# Increased fronto-temporal connectivity by modified melody in real music

**DOI:** 10.1101/2020.01.12.903609

**Authors:** Chan Hee Kim, Jaeho Seol, Seung-Hyun Jin, June Sic Kim, Youn Kim, Suk Won Yi, Chun Kee Chung

## Abstract

In real music, the original melody may appear intact, with little elaboration only, or significantly modified. Since a melody is most easily perceived in music, hearing significantly modified melody may change a brain connectivity. Mozart KV 265 is comprised of a theme with an original melody of “Twinkle Twinkle Little Star” and its significant variations. We studied whether effective connectivity changes with significantly modified melody, between bilateral inferior frontal gyri (IFGs) and Heschl’s gyri (HGs) using magnetoencephalography (MEG). Among the 12 connectivities, the connectivity from the left IFG to the right HG was consistently increased with significantly modified melody compared to the original melody in 2 separate sets of the same rhythmic pattern with different melody (*p* = 0.005 and 0.034, Bonferroni corrected). Our findings show that the modification of an original melody in a real music changes the brain connectivity.

## Introduction

Melody is a feature effortlessly extracted from music, which sometimes represents the music itself. Beethoven symphony No. 9 is known as “Ode to joy”, since the melody of “Ode to joy” appears repeatedly across the 4^th^ movement of the symphony, which is easily and prominently detected. It would be interesting to see how the brain reacts to phrases that contain prominent information such as a well-known melody while listening to real music, or phrases that do not. If this is something that can be detected without specific musical expertise, it would be interesting to see how the non-expert brain will react.

Purpose of our study was to verify the changes in non-expert brain by the musics including a well-known melody and its modifications using Mozart Variations. Mozart 12 Variations KV 265 on “Ah! vous dirai-je Maman” of a French chanson is well-known as “Twinkle Twinkle Little Star” Variations, of an English lullaby. In some of the variations, the original melody appears intact or with little elaboration only. In others, it is significantly modified. We call the former variations with original melody (VOM) and the latter variations with modified melody (VMM). The original melody of “Twinkle Twinkle Little Star” in the *Theme* is modified or repeated in 12 variations. Five conditions of the *Theme, Variation I, Variation II, Variation III,* and *Variation IV* are composed in the same key, harmony, meter, timbre, and ternary form. Among the five conditions, the original melody is present in the *Variation II* and *IV.* However, the original melody is significantly modified in the *Variation I* and *III*. In terms of rhythmic pattern, the *Variation I* and *II* are of 16^th^ notes, while the *Variation III* and *IV* are of 8^th^ notes. Hence, two sets of *“Variation I* vs. *II*” and *“Variation III* vs. *IV”* are of the same rhythmic patterns, but of different melodies (Fig. 1).

**Fig 1.**
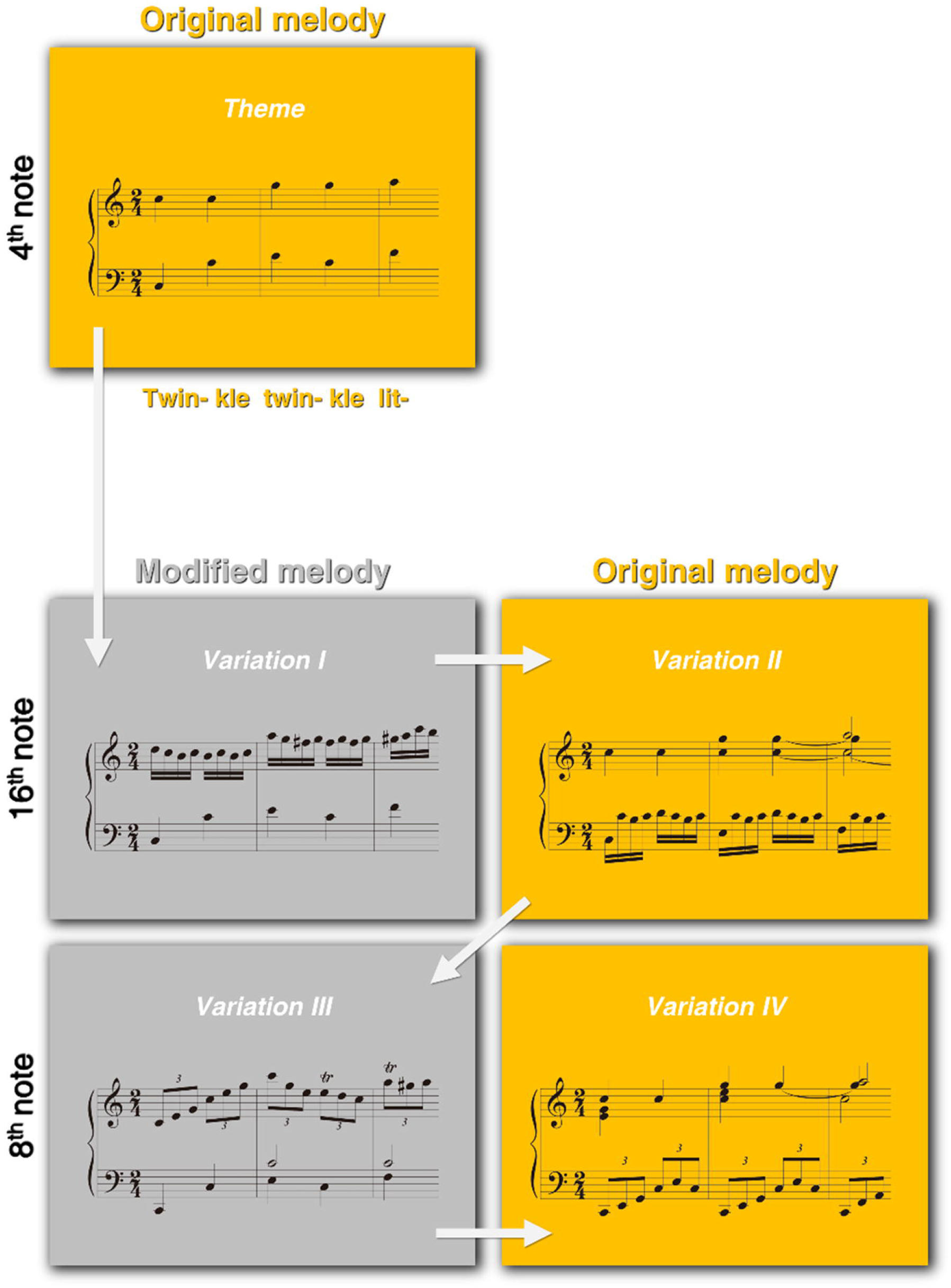
Musical stimuli. Five conditions of the *Theme, Variation I, II, III,* and *IV* selected in Mozart 12 Variations on “Ah! vous dirai-je Maman”KV 265. The original melody of “C5 - C5 - G5 - G5 - A5 (Twin- kle twin- kle lit-)” is included in the *Theme, Variation II,* and *Variation IV* (orange boxes). The *Variation I* and *Variation III* include the modified melodies (gray boxes). Rhythmic patterns are consistent in each set of *“Variation I* and *Variation II”* (16^th^ notes) and *“Variation III* and *Variation IV*” (8^th^ notes).

The human brain has several interconnected key regions processing music, including bilateral inferior frontal gyri (IFGs) and Heschl’s gyri (HGs) [1–8]. Each region and hemispheres are known to have a specific function. The right IFG is dominant in the processing of musical syntax, compared to the left IFG [1]. The left IFG is activated by pattern deviations of melodic contours [9], and is involved in the processing of familiarity in music [10]. The bilateral HGs are involved in pitch and melody process, which is dominant in right hemisphere [3, 5, 11]. The right HG is associated with the processing of auditory stream [12], a deviance of tone [13, 14] and spectral pitch [5]. Extracting and interpreting a melody in music would be accompanied with changes in a regional connectivity. A hemispheric or interregional connectivity between related regions in bilateral IFGs and HGs would allow to infer what characteristics the difference between VOM and VMM reflect.

Effective connectivity for 12 connections between the bilateral IFGs and HGs was calculated by linearized time delayed mutual information (LTDMI) [15]. LTDMI is a measure of functional coupling estimating information transmission between two time series based on mutual information (MI). Directional information transmission calculated by time delayed mutual information (TDMI) detects linear and non-linear correlation between two time series.

Therefore, we hypothesized that directional information transmission between related regions associated with the processing of intrinsic differences implied in the VOM and VMM would be reflected in the LTDMI between the bilateral IFGs and HGs. We further hypothesized that the two sets of *“Variation I* vs. *II”* and *“Variation III* vs. *IV”,* with the original melody and modified melody, would consistently change effective connectivity between the bilateral IFGs and HGs.

## Materials and Methods

### Ethics Statement

The study was approved by the Institutional Review Board of the Clinical Research Institute, Seoul National University Hospital (IRB No. C-1003-015-311). All experiments were conducted in accordance with the ethical guidelines and regulations.

### Participants

Twenty-five participants (15 females, mean age, 26.8 ± 3.4 years) were all right-handed (mean Edinburgh Handedness coefficient, 95.7 ± 7.1). All participants had normal hearing, and had not received any formal musical training. Prior to the experiments, all the participants provided the informed consent in a written form.

### Musical stimuli

Out of the theme and the 12 Variations in Mozart’s Variation on “Ah! vous dirai-je Maman” KV 265, we selected the five conditions of *Theme, Variation I, II, III,* and *IV* in Fig. 1, composed in the same key (C major), harmony (tonic-tonic-subdominant-tonic...), meter (2/4), and structure (ternary form). In magnetoencephalography (MEG) experiment, all participants were asked to passively listen to the musical stimuli without to paying any attention to any specific musical feature, such as melody.

### Time window

The time window was selected based on the concept of the motif [16]. The ascending five quarter notes of “C5 - C5 - G5 - G5 - A5” (2,100 ms) of the time window were in the motif of “C5 - C5 - G5 - G5 - A5 - A5 - G5 - G5”. A previous behavioral study reported that differences in melody were detected by listening to a segment of three to six notes in a motif of an original melody [17]. Even though it usually takes about 400 ms before the brain response leads to a motor response after stimulus onset, 5 quarter notes were considered appropriate time widow. Also we thought it was reasonable to focus on the original melody, even considering that familiarity and congruence, etc., which might differ between the conditions; i.e. responses to familiar and congruent conditions were faster [18, 19]. For the five quarter notes of in the *Theme,* the number of 16^th^ notes or 8^th^ notes (in triplets) subdivided in accompaniments was the same between the *Variation I, II, III,* and *IV.*

The present musical stimuli were recorded music, so those were presented without a break between movements of a theme and variations. To select the proper time window, we had to consider all the subdivided notes included in a quarter note in each time window in all conditions. The time window of 2,100 ms was the most suitable duration because (1) 2,100 ms covered the onset of the first quarter note to the offset of the last quarter note in the time window; (2) the number of additional notes did not exceed a quarter note in all the conditions; and (3) fade-out times of the final note in the preceding condition did not overlap with the onset of the time window in the following condition. The same phrases of the time window were repeated four times in each condition. However, to rule out the effect of repetition [20], we analyzed only the “C5 - C5 - G5 - G5 - A5” in the opening in each condition.

### Data recording

The MEG signal was recorded with a sampling frequency of 1,000 Hz using a 0.1–200 Hz band pass filter in a magnetically shielded room with a 306-channel whole-head MEG System (Elekta Neuromag Vector View™, Helsinki, Finland). Electrooculograms (EOG) and electrocardiograms (ECG) were also simultaneously recorded to remove ocular and cardiac noise at a later time. We eliminated the environmental magnetic noise of raw MEG signals with the temporal signal space separation (tSSS) algorithm in MaxFilter 2.1.13 (Elekta Neuromag Oy, Helsinki, Finland) [21].

During the MEG recording, for about five minutes, all participants sat in a magnetically shielded room and passively listened to music (Mozart 12 Variations on “Ah! vous dirai-je Maman” KV 265, 2011, Brilliant classics, Dutch). Musical stimuli were presented binaurally through plastic tubal silicone earpieces, 50 cm in length using STIM2™ (Neuroscan, Charlotte, NC, USA) at 100 dB. Visual stimuli of a silent movie clip (Love Actually, 2003, Universal Pictures, USA) were presented on a screen to keep the participants awake [22, 23] while the musical stimuli were being presented. For MEG signals, EOG, ECG, muscle artifact was removed using the independent component analysis. Epochs were determined from −100 ms to 2,100 ms after the onset of each condition, and the baseline of each epoch was from −100 ms to 0 ms.

### Data analysis

The source coordinates for the ROIs of the bilateral HGs and IFGs were based on the standard Talairach coordinates; Heschl’s gyrus (transverse, BA 41, BA 42) and the inferior frontal gyrus (triangular part, BA 45); the x, y, and *z in* Talairach coordinates (millimeters) were −53.5, −30.5, and 12.6 in the left HG, 55.4, −30.5, and 12.6 in the right HG, −55.5, 11.7, and 20.6 in the left IFG, and 53.5, 12.7and 20.6 in the right IFG, respectively (Fig. 2). In Fig. 2, The Talairach coordinates were visualized using BrainNet Viewer (http://nitrc.org/projects/bnvZ). The signal for each regional source of 4 ROIs (bilateral HGs and IFGs) was extracted using a 14–30 Hz band-pass filter for each participant using BESA 5.1.8.10 (MEGIS Software GmbH, Gräfelfing, Germany), which was averaged at MATLAB 7.7.0.471 (Math Works Inc., Natick, MA, USA). Beta band rhythm of 14–30 Hz underlies the top-down control [24], reflects a deviant word processing [25], and is related to the semantic evaluation for the spoken word [26] and categorical perception [27]. Considering that our hypothesis is focused on separating VOM and VMM, with the original and modified melody, we chose the beta band rhythm as the target frequency.

**Fig 2.**
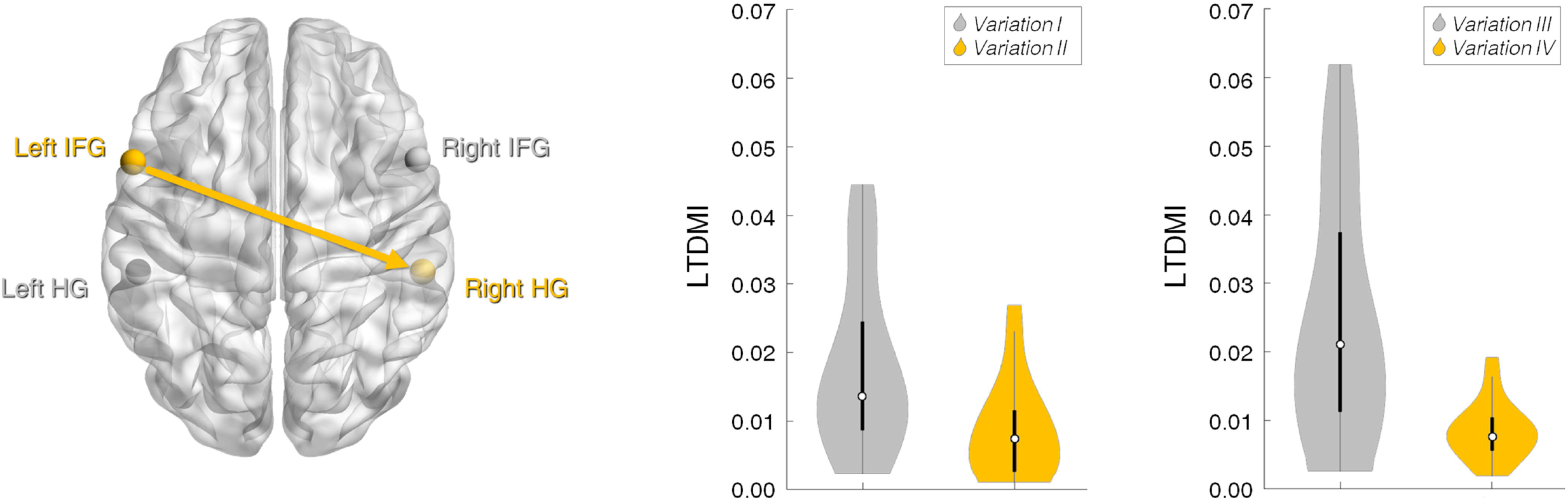
Difference in effective connectivity from the left IFG to the right HG for the modified and original melodies. Effective connectivity in both sets of *“Variation I* vs. *Variation II”* and *“Variation III* vs. *Variation IV”* showed an identical pattern in the Wilcoxon signed-rank test. Violin plots show that the LTDMI values are significantly higher for the *Variation I* and *III* with the modified melodies than for the *Variation II* and *IV* with the original melody; *“Variation I* vs. *Variation II”, Z* = −3.512, *p* = 0.005, Bonferroni corrected; *“Variation III* vs. *Variation IV”, Z* = −2.987, *p* = 0.034, Bonferroni corrected (See also Table 1).

For the twelve connections among the bilateral HGs and IFGs, effective connectivity for twelve connections was estimated using LTDMI [15]. The MI is a measure testing the relationship between the conditions by measuring the amount of information between two random variables. The LTDMI is an information theoretic measure estimating the directional information transmission of functional coupling based on MI:

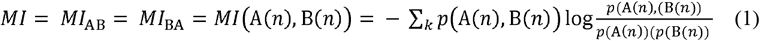

In case of *A*(*n*) = *B*(*n*), the MI is maximum. When two time series are independent, the MI is zero. The directional information transmission between the two time series estimated based on TDMI, time delayed mutual information [15]:

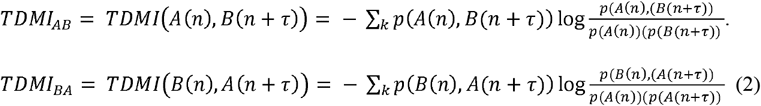

TDMI is asymmetric MI obtained by adding a time delay. Both of linear and non-linear correlation between two time series can be detected by TDMI. The data length of 400 ms epoch in our study was insufficient in reconstruction of a reliable PDF for general TDMI. Based on TDMI, the linear correlation for the information transmission between time series A and B is estimated by LTDMI:

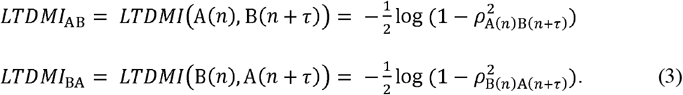

where τ is delay time, and *ρ*_A(*n*)B(*n*+*τ*)_ or *ρ*_B(*n*)A(*n*+*τ*)_ is a cross-correlation coefficient. The LTDMI value was averaged over delay time, which was calculated for every individual participant for each of the four conditions and twelve connections. See our previous study^3^ for the details of the LTDMI.

The mean LTDMI value for the time window of 2,100 ms in each condition was calculated for the twelve connections between the four ROIs. Because the data set was not normally distributed, the differences in LTDMI values of the conditions including “Twinkle Twinkle Little Star” melody or not were tested by the Wilcoxon signed rank test using SPSS 21.0 software (IBM, Armonk, NY, USA). Multiple comparisons for the twelve connections in each set were adjusted by the Bonferroni test (*p* < 0.05).

## Results

The Wilcoxon signed-rank test revealed that the LTDMI from the left IFG to the right HG was significantly different between *“Variation I* vs. *Variation II”* (*Z* = −3.512, *p* = 0.005, Bonferroni corrected) as well as between *“Variation III* vs. *Variation IV”* (*Z* = −2.987, *p* = 0.034, Bonferroni corrected) (Table 1). In this fronto-temporal connection, the LTDMI values increased in the VMM (*Variation I* and *III*) compared to the VOM (*Variation II* and *IV*) (Fig. 2).

**Table 1.** LTDMI difference between the conditions with the modified or original melodies. For the both sets of *“Variation I* vs. *Variation II”* and *“Variation III* vs. *Variation IV”,* the LTDMI values differed in a connection from the left IFG to the right HG among twelve connections (Wilcoxon signed-rank test, N = 25, Bonferroni corrected \**p* < 0.05 and \*\**p* < 0.01).

|  | *Variation I vs. Variation II* |  |  | *Variation III vs. Variation IV* |  |  |
| --- | --- | --- | --- | --- | --- | --- |
|  | *Z* | *P*<sub>(uncorrected)</sub> | *P*<sub>(corrected)</sub> | *Z* | *P*<sub>(uncorrected)</sub> | *P*<sub>(corrected)</sub> |
| *Left HG → Right HG* | -0.565 | 0.572 | 1.0 | -1.090 | 0.276 | 1.0 |
| *Left HG → Left IFG* | -0.727 | 0.468 | 1.0 | -1.843 | 0.065 | 0.784 |
| *Left HG → Right IFG* | -0.148 | 0.882 | 1.0 | -0.229 | 0.819 | 1.0 |
| *Right HG → Left HG* | -1.493 | 0.135 | 1.0 | -0.175 | 0.861 | 1.0 |
| *Right HG → Left IFG* | -1.286 | 0.199 | 1.0 | -1.029 | 0.304 | 1.0 |
| *Right HG → Right IFG* | -1.305 | 0.192 | 1.0 | -0.148 | 0.882 | 1.0 |
| *Left IFG → Left HG* | -1.117 | 0.264 | 1.0 | -1.655 | 0.098 | 1.0 |
| *Left IFG → Right HG* | -3.512 | 0.0004 | **0.005 \*\*** | -2.987 | 0.0028 | **0.034 \*** |
| *Left IFG → Right IFG* | -0.875 | 0.382 | 1.0 | -1.238 | 0.216 | 1.0 |
| *Right IFG → Left HG* | -1.129 | 0.259 | 1.0 | -0.498 | 0.619 | 1.0 |
| *Right IFG → Right HG* | -1.762 | 0.078 | 0.936 | -0.982 | 0.326 | 1.0 |
| *Right IFG → Left IFG* | -2.193 | 0.028 | 0.340 | -0.848 | 0.397 | 1.0 |

## Discussion

Effective connectivity from the left IFG to the right HG increased when the original melody of “Twinkle Twinkle Little Star” was significantly modified (VMM), compared with that when the original melody was intact (VOM). It appeared consistent in two different sets with the same rhythmic patterns but with the different melody pattern.

The present study employed real music with its variations as musical stimuli, dissected musical elements in each condition, and devised the two comparable sets of conditions, which have the same rhythmic pattern but different melody. Here, we exploited naturalistic conditions in real music instead of devising artificial conditions, and successfully demonstrated how significant variations of melody in real music change a regional connectivity in the human brain. These are distinct from the attempts in previous studies used to original musical pieces [10, 28–31].

An fMRI study, using a real music of “Adios Nonino”, studied the effects of motif repetitions on effective connectivity changes. Specifically they found that the hippocampal connectivity is modulated by motif repetitions, showing strong connections with working memory-relevant areas [31]. This study also shows that a real music and characteristics inherent in it change the brain connectivities. However, unlike this study, our study did not only focus on the motif of monody within the music. Instead, we identified how the motif was intertwined with other elements of music through music analysis, and tried to present results that did not exclude all possible elements within the music.

When hearing real music the brain regions would interact to support complex cognitive processes [32, 33]. In the perspective of the brain networks, the change in effective connectivity from the left IFG to the right HG could be interpreted in the syntactic process. However, modified and original melody detection would be different processes from syntactic process, although relatively similar cortical regions may be involved [1, 23, 34]. The left IFG is highly activated for the detection of novelty within a musical context based on long term memory [35].

The left IFG is also involved in the retrieval of working memory [36] and episodic memory [30], and in the processing of familiarity [10]. Also, the left prefrontal area is activated for the recognition of pattern deviations in melody [9]. On the other hands, the right HG is dominant for the processing of stream segregation for auditory stimuli [12], and is highly sensitive for a deviance of tone [13, 14] and spectral pitch [5], compared to the left hemisphere. The right superior temporal sulcus is also involved in the processing of familiarity [37]. In this regards, increased and decreased fronto-temporal connectivity would involve memorizing melody or streams, feeling the familiarity, recognizing patterns of melody, and detecting tone deviation or novelty within the VMM with modified melody and the VOM with the original melody.

Also it is interesting that the change in fronto-temporal connectivity was observed in the beta frequency band (see also the Materials and Methods section). The connectivity changes were consistent in the two sets of *“Variation I* vs. *II”* and *“Variation III* vs. *IV”,* which are different in the rhythmic patterns of 16^th^ and 8^th^ notes sequences, respectively. Generally the beta frequency band is associated with the processing of deviant stimuli [25] and memory/categorical perception [27]. The connectivity in the beta frequency band would reflect the categorization of the deviant VMM and the VOM based on the information of the original melody acquired in real-life.

Also, beta frequency is related with beat perception. While regular auditory stimuli were presented, beta oscillation was synchronized with the beat, while it was increased when the beat was omitted [38, 39]. Therefore, the changes in fronto-temporal connectivity at the beta frequency band might reflect omission of the beats on the original melody of quarter notes sequence in each condition, regardless of changes in rhythmic patterns. It may be an alternative explanation why the fronto-temporal connectivity increased with modified melody conditions compared to the original melody.

In our study, the time window of five notes, “C5 - C5 - G5 - G5 - A5” (2.1 sec) was set based on the “C5 - C5 - G5 - G5 - A5 - A5 - G5 - G5” motif of “Twinkle Twinkle Little Star” song in the *Theme.* In a previous study, the isolation point, the point at which a melody is correctly identified, is about 2.5 sec of 5 notes for highly familiar melodies [17]. Therefore, the five notes in the motif, which we analyzed were long enough to detect the original melody.

Connectivity results in the two sets of conditions, with different rhythms, showed similar pattern. The hypothesis of our study was to find a connection that connectivity pattern was consistent for the two sets of “VOM and VMM” of the different rhythmic pattern. Therefore, the results would be by pitch rather than rhythm. Therefore, it would be natural that rhythmic affects are excluded from our results. Furthermore, the one that influenced the results could be eventually be dismissed as “original melody and modified melody” that encompasses all of familiarity, the stimuli characteristics, and the variation identification.

There are several limitations in our study. The present study was limited to 4 regions of interest, including bilateral inferior frontal and superior temporal area. There should be further studies covering the whole brain. For current findings targeting non-musicians, comparative studies involving musicians are needed. Finally, in order to focus on melody difference, we tried our best to eliminate the effects of other elements implied in naturalistic conditions. However, it should be discovered through further studies.

In conclusion, the fronto-temporal connectivity from the left IFG to the right HG was enhanced when the modified melody was presented, compared with the original melody of “Twinkle Twinkle Little Star” in Mozart “Ah! vous dirai-je Maman” KV 265. Our findings show that the modification of an original melody in a real music changes the brain connectivity.

## Acknowledgements

We would like to thank Ji Hyang Nam for the technical support in MEG acquisition.

## Notes

### Competing Interest Statement

The authors have declared no competing interest.

